# Comparative analysis of iPSC-derived human kidney organoid and tubuloid models

**DOI:** 10.1101/2024.11.24.625103

**Authors:** Ana B. Nunez-Nescolarde, Leilani L. Santos, Lingyun Kong, Elif Ekinci, Paddy Moore, Evdokia Dimitriadis, David J. Nikolic-Paterson, Alexander N. Combes

## Abstract

**Background:** Epithelial kidney organoids (tubuloids) made from kidney biopsies, urine, or iPSC-derived kidney organoids offer new opportunities in experimental and clinical nephrology. Yet, we have limited knowledge of how tubuloid models differ from each other, from iPSC-derived kidney organoids and from the human kidney. New insight is required to guide model selection for studies in kidney physiology and disease.

**Methods:** Tubuloids were generated from adult nephrectomy samples (adult tubuloids n=3), iPSC-derived kidney organoids (iTubuloids n=3), and for the first time, from human fetal kidneys (fetal tubuloids n=3). Kidney organoid and tubuloid models were compared to each other and to adult human kidney using bulk RNA sequencing. Tubuloids were subject to hypoxic insults over three passages and profiled with RNA sequencing to assess cumulative injury and repair.

**Results:** Expression signatures of proximal and distal tubules were stronger in adult kidneys than any organoid or tubuloid model. Comparative analysis between the models revealed that iPSC-derived kidney organoids had the highest expression of proximal tubule markers, even though adult tubuloids were derived from mature kidneys. Collecting duct signatures were enriched in adult and fetal tubuloids. Adult tubuloids showed stronger signatures of ageing and inflammation, while fetal tubuloids had enhanced ureteric tip progenitor signatures. Over 80 genes linked to inherited disorders were expressed in all tubuloid cultures, while a further 54 were expressed at higher levels in either adult, fetal or iTubuloids. Tubuloids subject to a single hypoxic injury effectively recovered by the end of the passage, while iTubuloids exposed to hypoxia over three passages expressed markers of maladaptive repair.

**Conclusions:** This study provides new transcriptome-wide reference data to aid in the selection and optimization of disease modelling for the human kidney. It defines common and unique opportunities to model inherited disorders in adult, fetal and iTubuloid models and illustrates new potential to model repetitive injury in long-lived tubuloid cultures.

## INTRODUCTION

During organogenesis, an average of one million nephrons develop in each human kidney. Nephrons consist of a glomerulus – the filtering unit – and the tubular epithelium. The renal tubular epithelium plays a central role in the fluid and solute handling functions of the kidney and is vulnerable to a variety of genetic, environmental and metabolic disorders (1). While animal models have advanced insight into kidney disease and driven target discovery, constraints on throughput and translational challenges are known limitations (2). Similarly, monolayer cultures of human kidney cell types lack the expression profiles, interactions and structure of nephrons. A such, human kidney organoids provide a promising alternative to advance our understanding and treatment of kidney disease, with demonstrated applications in acute kidney injury, inherited disorders, and infection (3–8).

Induced pluripotent stem cells (iPSCs) derived from adult somatic cell types are a favored starting point for kidney organoids as they are widely available and amenable to genetic manipulation. Most protocols involve modulating developmental signalling pathways to promote formation of either nephron or collecting duct progenitor cell types (9–14). Differentiation protocols targeting nephron formation contain stromal cells, endothelial cells and nephron tubules that are appropriately segmented into proximal, medial, and distal identities (15, 16). In most protocols, epithelial cell types take time to form and only remain viable and healthy for limited time (16–18), though extended cultures have been reported (19, 20).

An alternative long-term adult-derived kidney organoid model has been developed by culturing cells derived from a nephrectomy or urine samples in R-Spondin-conditioned medium containing FGF10, which promotes the survival and proliferation of epithelial progenitor cells (21–23). Renal epithelial cells cultured in this medium form cystic ‘tubuloid’ structures, each expressing markers of a single cell type including proximal tubule, loop of Henle, distal tubule and collecting duct (21). Cryopreserving and passaging tubuloid lines distinguishes them from iPSC-derived kidney organoids, which are typically generated per experiment, though cryopreservation of monolayers and organoids has been reported (24, 25). The production of iPSC organoid-derived tubuloids (originally iPSCodTubuloids, abbreviated here as iTubuloids) is also possible using adult-derived tubuloid culture conditions (26). However, it remains unclear how well adult-derived tubuloids and iTubuloids model the expression patterns of the cell types they represent, or how they compare to iPSC-derived kidney organoids, particularly regarding mature nephron signatures. In this study, we made tubuloids from iPSC-derived kidney organoids, adult nephrectomies, and the developing human kidney, comparing these to cortical nephrectomy samples and iPSC-derived kidney organoids using bulk RNA sequencing (RNAseq). Analysis of cell type and maturation signatures contrasts with prior reports and provides a valuable resource to evaluate the suitability of tubuloid cultures for specific disease modelling applications. In addition, we explore new possibilities to model cumulative injury by identifying signatures of maladaptive repair in iTubuloids exposed to multiple periods of acute hypoxia.

## METHODS

### Generation of iPSC-derived kidney organoids and adult-derived tubuloids

iPSC-derived kidney organoids were generated using a modified version of the Takasato protocol (9) as previously described (27). Adult tubuloids were generated from three nephrectomies as per Schutgens et al., (21, 22), with the exception that the dissociation of renal epithelial fragments was performed at 37°C in a gentle shaker.

### Establishment of iTubuloids

A protocol to derive tubuloids from iPSC-derived kidney organoids has recently been published (26). Our method was developed independently and is outlined as follows. d20 iPSC-derived kidney organoids (∼2 million cells) were digested in 1 ml TrypLE Select Enzyme (1X) for 10–12 min at 37 °C. Samples were centrifuged (400 x g, 10 min, 4 °C), resuspended in Basic Medium (ADMEM/F12, 1% Glutamax, 1% HEPES and 1% Pen/Strep), filtered through a 70 μm strainer, and centrifuged again (400 x g, 7 min, 4 °C). The pellet was resuspended in 40 μl ice-cold Matrigel and plated in a prewarmed 24-well suspension plate. After 20 min at 37 °C, 500 μl warm Expansion Medium (ADMEM/F12, 1% Glutamax, 1% HEPES, 1% PEN/Strep, 1.6% B27 supplement, 1% R-spondin-3-conditioned medium, 50 ng ml−1 EGF, 100 ng ml–1 FGF-10, 1 mM N-acetyl-L-cysteine, 10 µM-27632, 5 μM A8301 and 0.1 mg ml–1 primocin) was added. iTubuloids were cultured at 37 °C, 5% CO2, and 19% O2, with medium changes every 2–3 days. Epithelial structures formed within 2–3 days and were passaged weekly. Multiple iPSC-derived kidney organoids, from three different genetic backgrounds (including gene-edited iPSCs), all yielded iTubuloid cultures within days.

### Establishment of fetal tubuloids

Half of a 12–14-week gestation human fetal kidney was used to seed one well of fetal tubuloids in a 24-well plate. On collection, tissue can be stored in Basic Medium at 4 °C for up to 24 h before processing. Samples were minced with sterile scalpels and transferred to a 15-ml falcon tube using 5 ml cold PBS. Tissue fragments were pipetted with a 10 ml serological pipette, allowed to settle, and the excess PBS, along with floating fat and debris, was removed. This was repeated three times using cold PBS and a 5 ml pipette to further break the tissue. The fragments were then incubated in 1 ml prewarmed TrypLE Select Enzyme (1X) for ∼20 min at 37 °C, with pipetting every 5 min to aid dissociation. After centrifugation (400 x g, 7 min, 4 °C), cells were resuspended in Basic Medium. The protocol proceeds as described above for iTubuloids with fetal forming within 2–3 days and passaged weekly.

### Passage of tubuloids

Tubuloids were passaged at a 1:2–1:3 ratio. Ice-cold DMEM (1–3 ml) was added to detach tubuloids, which were then dissociated by pipetting (∼30 times with a P1000) and centrifuged (400 x g, 7 min, 4 °C). The pellet was resuspended in 1 ml ice-cold Basic Medium, transferred to a microtube, and further dissociated using a P10 tip attached to a P1000. After a second centrifugation (400 x g, 10 min, 4 °C), the pellet was resuspended in ice-cold Matrigel. A 30–40 μl aliquot was used to seed each new well in a 24-well plate.

### RNAseq and analysis

Total RNA was isolated using the Isolate II RNA Mini Kit (Bioline, BIO-52072). Sequencing was performed at MHTP Medical Genomics facility using a 3’-Multiplex RNA-seq method that captures and sequences the 3’ end of polyadenylated transcripts (28). All samples were processed starting with 25 ng of total RNA on an Illumina NextSeq 2000, which resulted in a loading metric of 99.7 and 90% >Q30. 463.2 million reads were assigned to indexes.

Fastq files were processed using the Laxy tool (29) and the NfCore/RNAseq (v3.10) pipeline, using the--with_umi function (30). Reads were aligned to the ENSMBL Homo sapiens GRCh38 reference (release 109) using STAR aligner and quantified using Salmon producing the raw genes count matrix (31). Various quality control metrics were generated and summarised in a multiQC report (32). Raw counts were then analysed with Degust (33), a web tool that performs normalization using trimmed mean of M values (TMM) (34), and differential expression analysis using limma/voom (35). Heatmaps were generated using mean-centering data on the varistran R package (36). Differentially expressed genes (adjusted p < 0.05) were used to perform enrichment analysis with DAVID (37) and plotted using GraphPad prism v9. Venn diagrams created with Venny v2. Threshold for volcano plots were log2FC > 1 and FDR < 0.001.

See Supplementary Methods for further details on methods and catalogue numbers for key consumables and reagents (Supplemental Table 1).

## RESULTS

### Establishment of tubuloids from nephrectomies, iPSC-derived kidney organoids and human fetal kidney

Tubuloids have been previously generated from adult nephrectomy samples, cells isolated from the urine, and recently from iPSC-derived kidney organoids (21, 26). Here, tubuloids were established from three adult nephrectomies (adult tubuloids), and three batches of iPSC-derived kidney organoids, each from a unique iPSC line (Figure 1A). Tubuloids from iPSC-derived kidney organoids (iTubuloids) started to form within 2-3 days and grew to resemble adult tubuloids (Supplemental Videos 1-2).

**Figure 1.**
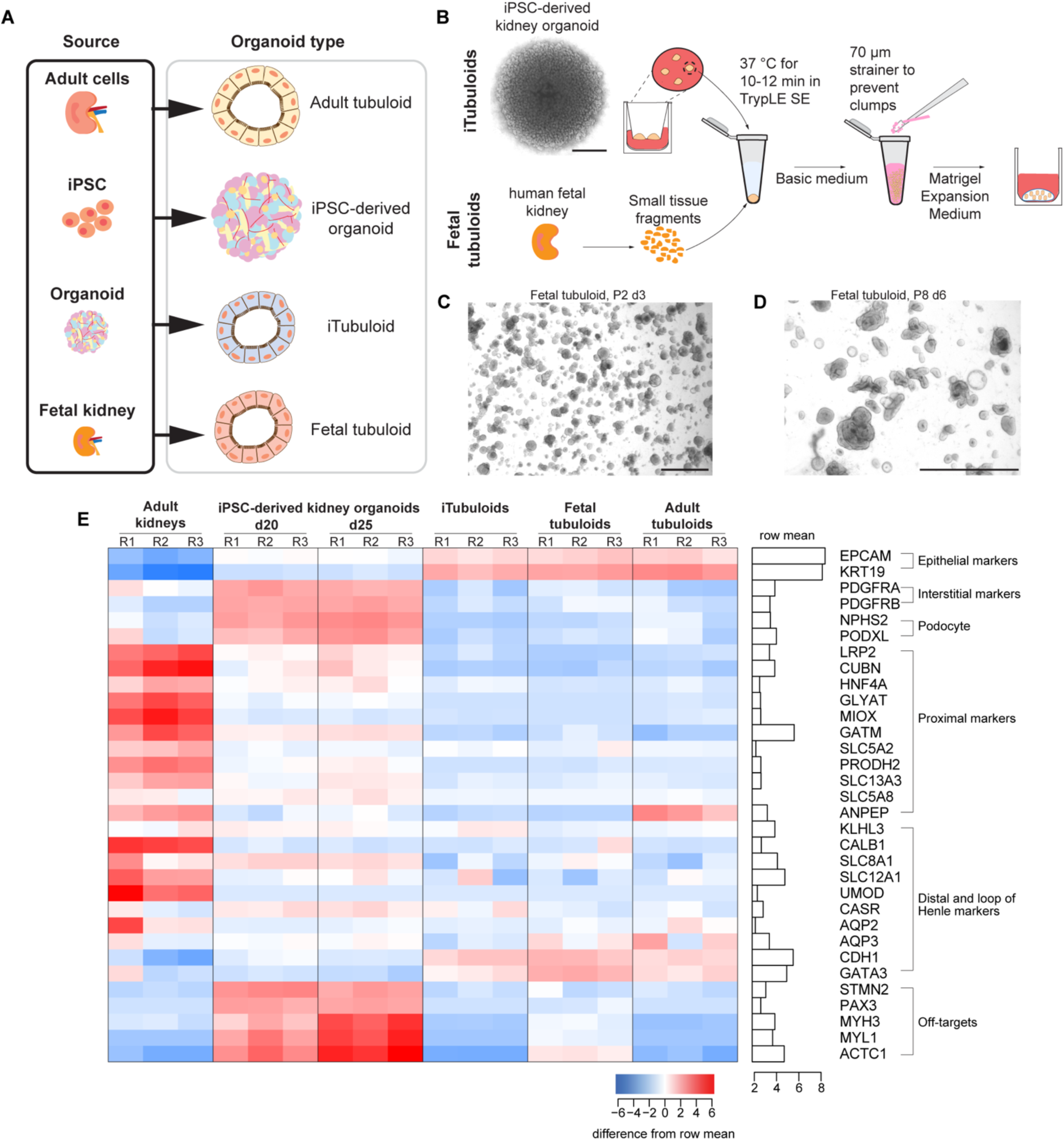
Comparison of human kidney organoids and tubuloids with the human kidney. (A) Illustration of different types of kidney organoids analyzed in this study. (B) Overview of iTubuloid and fetal tubuloid generation. (C-D) Representative bright field images of fetal tubuloids at day three (d3) after the second passage (P2), and d6 after P8. Scale bars = 1000 μm. (E) Heatmap of Log2 CPM-normalized expression levels of three biological replicates of human kidney tissue samples (two XY, one XX), d20 and d25 iPSC-derived kidney organoids (three XX), iTubuloid lines (two XX, one XY), human fetal tubuloids (three XX) and three human nephrectomy-derived tubuloids (two XY, one XX).

In addition, we report for the first time, the generation of tubuloids from three human fetal kidneys aged 12-14 weeks of gestation (fetal tubuloids) (Figure 1A-B). 3D structures were visible within 2 days of seeding and maintained in culture over multiple passages (Figure 1C-D). Growth rates and recovery from cryopreservation were similar in tubuloids derived from adult nephrectomies, iPSC-derived kidney organoids and fetal kidney tissue. Likewise, tubuloid numbers decreased and fibroblast-like cells increased in cultures that were passaged >8 times from all starting material (Supplemental Figure 1).

### Comparative analysis of kidney organoids and tubuloids with the human kidney

The expression profiles and relative maturation of cell types within adult, fetal and iTubuloid cultures remains unclear and has a significant bearing on their utility for disease modelling. To investigate these questions, we performed bulk RNAseq on 1) iPSC-derived kidney organoids at day 20 and day 25 of differentiation, 2) three iTubuloid lines, 3) three fetal tubuloid lines and 4) three adult tubuloid lines, and 5) three human cortical kidney samples as a primary tissue reference.

We first compared markers of major cell types between iPSC-derived organoids, tubuloids and the adult kidney. The epithelial nature of tubuloid cultures was confirmed by increased expression of *EPCAM* and *KRT19* across all tubuloid types compared to iPSC-derived kidney organoids and cortical kidney samples (Figure 1E). As expected, iPSC-derived kidney organoids retained stromal/interstitial cell types (*PDGFRA, PDGFRB*) and podocyte signatures (*NPHS2, PODXL),* which were depleted in tubuloid cultures (Figure 1E). Select markers of the proximal tubule, distal tubule, and loop of Henle were strongly expressed in the cortical kidney samples, and at lower levels in *the invitro* models (Figure 1E). Expression of the loop of Henle marker *UMOD* was not detected in iPSC-derived kidney organoids or any tubuloid model. Broad markers of the distal tubule and collecting duct (*GATA3, CDH1)* were found at similar levels in all tubuloid groups (Figure 1E). Consistent with findings by Yousef Yengej et al. (26), we observed that markers of off-target muscle (*ACTC1*, *MYL1, MYH3)* and neural (*PAX3, STMN2*) cell types in iPSC-derived kidney organoids diminished upon iTubuloid derivation (Figure 1E).

### Comparison of epithelial cell signatures in organoid and tubuloid models

Tubuloid models established from iPSC-derived kidney organoids, human fetal kidneys and adult nephrectomies all displayed a mix of cystic and convoluted morphologies (Figure 2A). To gain insight into how these *in vitro* models compare on the cellular and molecular level, we evaluated their expression profiles using a multidimensional scaling plot. Clustering of samples within each experimental group suggests similarities between biological replicates, with adult tubuloid cultures showing the highest within-group variation. The three tubuloid types were more similar to each other than to iPSC-derived kidney organoids, with fetal and adult-derived tubuloids clustering in close proximity (Figure 2B).

**Figure 2.**
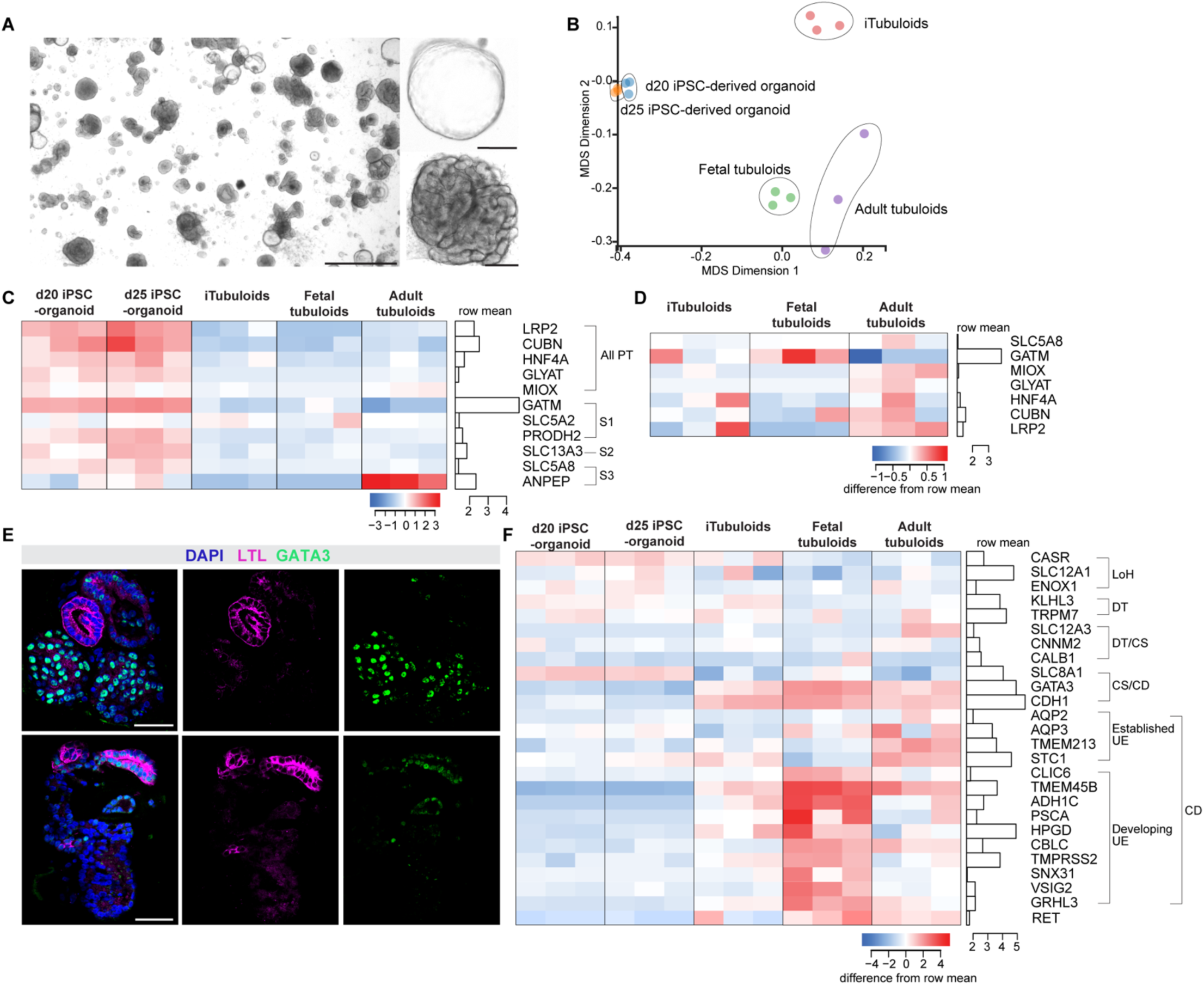
Nephron and collecting duct signatures vary between tubuloids and iPSC-derived kidney organoids. (A) Representative image of cystic and convoluted morphologies observed in all tubuloid cultures. Scale bars: left 1000 μm, right 200 μm. (B) Multidimensional scaling plot (MDS) plot of three biological replicates for d20 and d25 iPSC-derived kidney organoids, iTubuloids, adult tubuloids and fetal tubuloids. (C) Heatmap of Log2 CPM-normalized expression of general and segment 1-3-enriched proximal tubule markers. (D) Heatmap of Log2 CPM-normalized expression levels of mature proximal tubule markers in tubuloids. (E) Fetal tubuloids stained with LTL (proximal tubule marker) and GATA3 (distal marker). Scale bars = 50 μm. (F) Heatmap of Log2 CPM-normalized expression levels of distal nephron and loop of Henle markers.

iPSC-derived kidney organoids are known to resemble the first trimester of kidney fetal development (38), whereas adult-derived tubuloids could represent more mature cell types. Despite their immature nature, iPSC-derived kidney organoids expressed proximal tubule markers like *LRP2, HNF4A* and *CUBN* at higher levels than all three tubuloid models, with the exception of ANPEP, which is also expressed in the connecting tubule and the collecting duct (39) (Figure 2C). Most markers enriched in proximal tubule segments 1-3 in mature kidneys were not maintained in adult tubuloids, though some were expressed in iPSC-derived kidney organoids (Figure 2C). Amongst the tubuloid models, proximal tubule markers were expressed at higher levels in adult tubuloids (Figure 2D). However, all cultures contained tubuloids labelled with proximal tubule marker LTL and the distal marker GATA3 (Figure 2E). In some cases, these markers overlapped (Figure 2E), which is not observed in the developing human kidney (Supplemental Figure 2).

We then assessed key markers of the distal nephron and loop of Henle across the organoid models. All showed comparable expression levels of broad loop of Henle markers (*SLC12A1, CASR, ENOX1*) as well as markers for the distal tubule and connecting segment (*KLHL3, CNNM2, SLC12A3*) (Figure 2F).

Collecting duct signatures were not expected in our nephron-focused iPSC-derived kidney organoids, or the iTubuloids derived from these organoids. Markers upregulated in the established collecting duct/ureteric epithelium (*AQP2, AQP3, TMEM213, STC1*) were robustly expressed in adult tubuloids, but not in the other models. Markers enriched in the developing ureteric epithelium (*PSCA, HPGD, CBLC, RET, ADH1C, SNX31)* were increased in fetal tubuloids (Figure 2F) (40–42). Lower levels of ureteric epithelium markers in iTubuloids likely reflect the strong overlap in expression signatures of the distal nephron and collecting duct (38).

### Comparison of nephron development, repair, and ageing signatures

Nephron formation in iPSC-derived kidney organoids models the mesenchyme to epithelial transition that occurs during kidney development, whereas tubuloid formation is more akin to epithelial repair. The extent of mechanistic conservation between nephron development and repair remains unclear, as does whether a specific progenitor population, or any mature cell can repair an injured nephron. In agreement with previous studies (5, 43), markers of mesenchymal nephron progenitor cells (*CITED1, SIX2, GDNF*) were expressed in iPSC-derived kidney organoids and absent in all three tubuloid models (Figure 3A). Markers of the developing nephron (*JAG1, LHX1, EMX2*) were expressed in iTubuloids, fetal tubuloids and adult tubuloids (Figure 3A), though each gene is also expressed in adult nephron and collecting duct cell types. Similar levels of proliferative markers (*PCNA, MKI67*) were observed in the different organoid models; however, adult tubuloids expressed the highest levels of genes associated with glycolysis and fatty acid oxidation, which might indicate improved cellular maturation and greater energy consumption (Figure 3B-C). Markers associated with nephron repair and regeneration (*SOX9, NT5E, PROM1, CD44,* CD24) were detected in all three tubuloid types (highest in adult tubuloids), and absent in iPSC-derived kidney organoids (Figure 3B-C). Genes associated with aging (*UPP1, MLKL, CDKN2A, OSMR, B2M)* and inflammation (*LCN2, STAT3, JUN, CXCL1, ICAM1, SAA1)* (44–46) were expressed at higher levels in adult tubuloids compared to any other tubuloid model or iPSC-derived kidney organoids, which might reflect the effect of ageing *in vivo* (Figure 3D).

**Figure 3.**
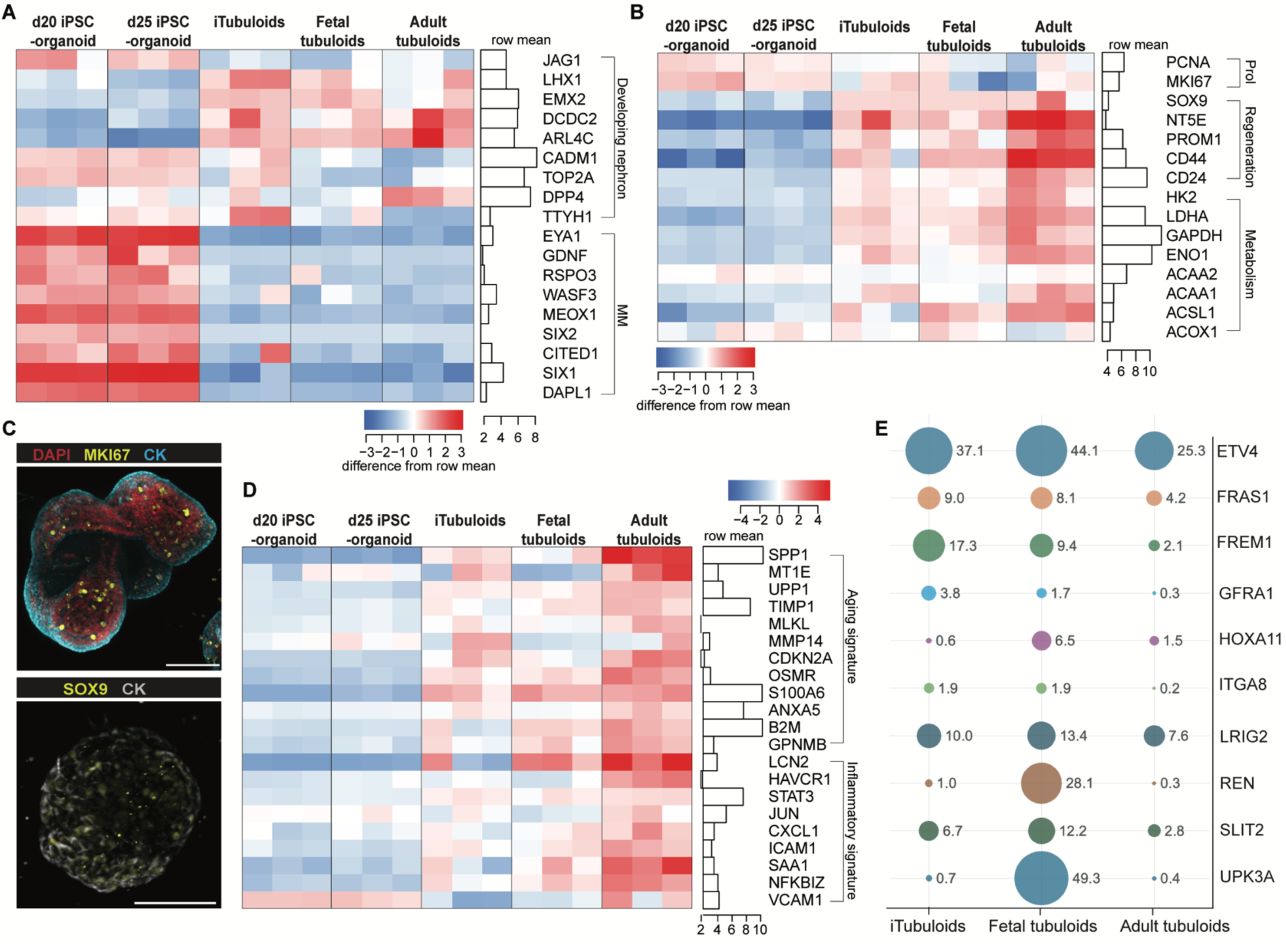
Signatures of development, repair, ageing, and inherited disease. (A) Heatmap of Log2 CPM-normalized expression levels of markers of peritubular aggregates (PTA) and renal vesicles (RV) as well as metanephric mesenchyme (MM) in the different organoid models. (B) Heatmap of Log2 CPM-normalized expression levels of markers of proliferation (Prol), regeneration and metabolism. (C) Representative immunofluorescence images of adult tubuloids showing the epithelial marker cytokeratin (CK), regeneration/repair marker SOX9 and proliferation marker MKI67. Both scale bars 100 μm. (D) Heatmap of Log2 CPM-normalized expression levels of markers related to aging and inflammation. (E) Bubble plot showing average CPM levels of genes representing monogenic causes of human CAKUT in iTubuloids, foetal tubuloids and adult tubuloids. Bubble size reflects CPM values (displayed on the right-side) and bubble color is unique to each gene (represented in y-axis).

### Potential to model inherited disorders

The capacity to derive tubuloids from adult and fetal kidney provides new possibilities to model inherited disorders. To explore this potential, we assessed the expression of 460 genes associated with inherited kidney disease in iTubuloids, fetal, and adult tubuloids. Unique gene symbols from the Kidneyome_SuperPanel (Version 8.79, https://panelapp.agha.umccr.org/panels/275/) were cross-referenced to single cell RNAseq data from the developing mouse and human kidney (13, 47) to identify 187 genes expressed in renal epithelial cell types in vivo. 134 out of the 187 epithelial genes were expressed in at least one tubuloid model (average CPM >5 from the three biological replicates). Disease-associated genes expressed in tubuloids included those normally expressed in the proximal and distal tubule, and ureteric epithelium (Supplemental File 1). Although most of these genes were expressed across all tubuloid cultures, a two-way ANOVA with Tukey’s multiple comparisons test identified 55 genes including *RET, UPK3A, SCNN1B, BMP7, FXYD2, SLC3A1* that were specific to, or enriched in, one or more tubuloid type (Supplemental File 1).

To date, mutations in 54 genes have been associated with congenital anomalies of the kidney and urinary tract (CAKUT), contributing to the development of chronic kidney disease (CKD) in patients under 30 years of age (48). Several of these genes were expressed at higher levels in fetal tubuloids compared to adult (Figure 3E). Interestingly, iTubuloids displayed gene expression levels intermediate between those observed in fetal and adult tubuloids. Altogether, these data define broad potential to study inherited disorders in tubuloids and highlight genes that may benefit from modelling in fetal, adult, or iTubuloid cultures.

### Modelling repetitive injury in iTubuloids

Up to one third of patients hospitalized with acute kidney injury (AKI) are at risk of rehospitalization within 12 months of discharge due to recurrent AKI events, increased risk of developing chronic kidney disease (CKD) and progression to kidney failure (49). The ability to passage tubuloids offers a unique platform to study regeneration and repair mechanisms following repeated injury. We explored this potential using hypoxia to model ischemic injury, which is a common clinical driver of AKI (50). iTubuloids were divided in three groups: control, one hypoxic exposure, and three hypoxic exposures (Figure 4A). Cultures were maintained over 3 passages for a total of 33 days (33 d). The control group was maintained in normoxic (19% O_­_) conditions for the duration of the experiment. Hypoxic exposures involved culture in hypoxia (1% O_­_) from days 4-6 after passage, followed by 5 days in normoxic conditions to allow repair (harvest d11) (Figure 4A).

**Figure 4.**
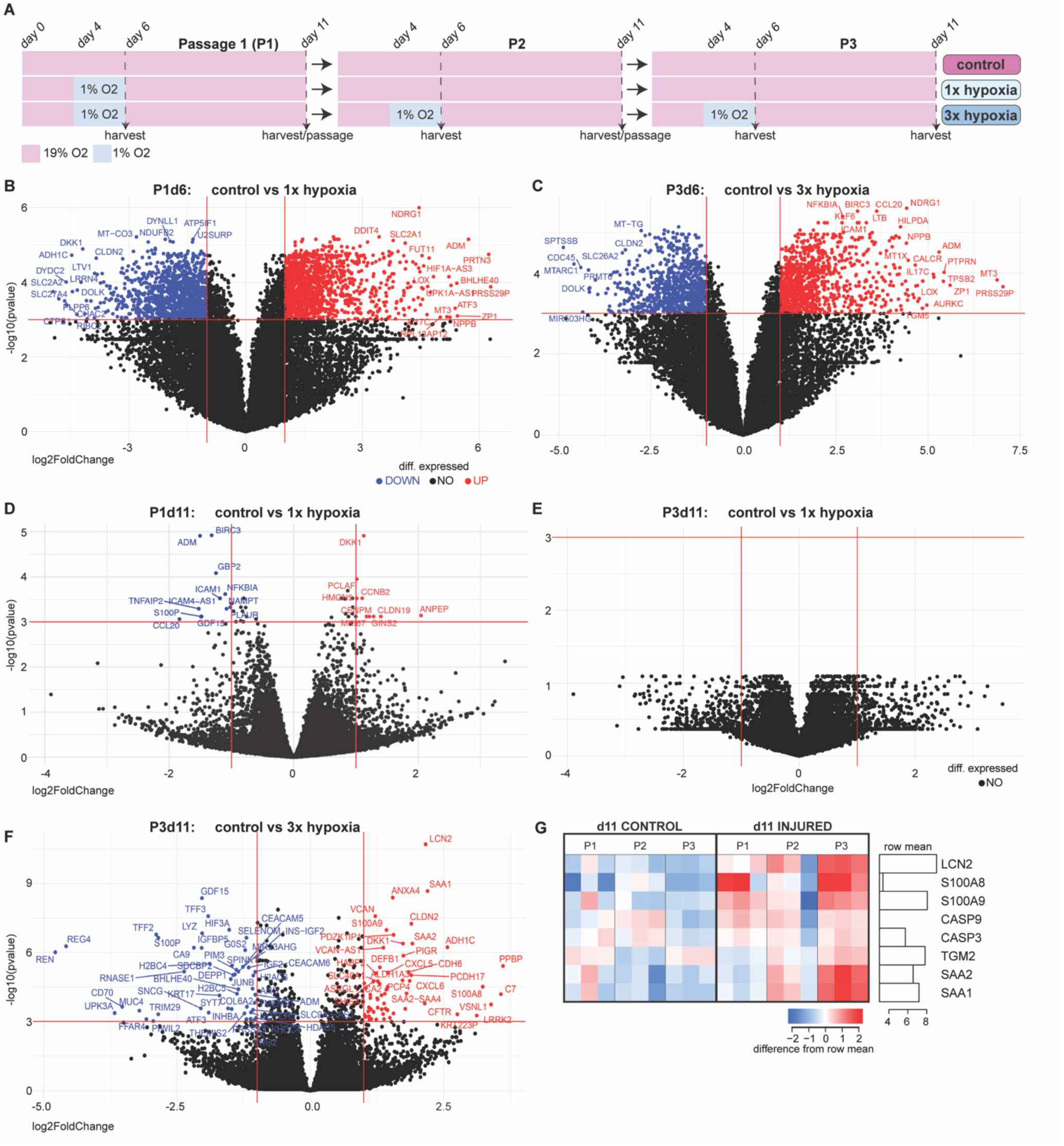
Repetitive hypoxic injury triggers signatures of maladaptive repair in iTubuloids. (A) Overview of the hypoxic injury experiment across three passages (P), in which iTubuloids are divided in three groups: control, 1x hypoxia and 3x hypoxia and collected for RNAseq. (B-F) Volcano plots of differentially expressed genes (log2 Fold Change > 1 and adjusted p-value < 0.001). Blue indicates downregulated genes; red, upregulated; no significant change in black. (B) day 6 3x hypoxia vs control (P3 d6). (D) Day 11 passage 1 1x hypoxia vs control (P1 d11). (E) Passage 1 day 6 1x hypoxia vs control (P1d6). (C) Passage 3 Passage 3 day 11 1x hypoxia vs control (P3 d11). (F) Passage 3 day 11 3x hypoxia vs control (P3 d11). (G) Heatmap of maladaptive repair markers (Log2 CPM-normalized expression) in three biological replicates for day 11 control and injured iTubuloids at P1, P2 and P3.

KEGG pathway analysis of iTubuloids exposed to hypoxia after one passage (1x hypoxia; assessed on P1 d6) revealed upregulation of signatures related to hypoxic injury including HIF signalling, metabolism, and inflammation (Supplemental Figure 3A). Similar pathways were observed in iTubuloids exposed to hypoxia after three consecutive passages (3x hypoxia; P3 d6), with 75% of genes upregulated in both 1x and 3x hypoxia groups (Supplemental Figure 3A-C). Genes associated with hypoxia (*NDRG1, BIRC3*) and acute kidney injury (*CXCL1, GDF15, JUN, NFKBIA, ICAM1, IL17*) (51) were amongst the highest upregulated in 1x and 3x hypoxic groups (log_­_Fold Change > 1 and FDR < 0.001) (Figure 4B-C, Supplemental File 2). Despite the similarity in the response to 1x and 3x hypoxic injuries, repair signatures differed. After the recovery period, the 1x hypoxic group (P1 d11) exhibited minimal transcriptional differences to stage-matched controls, and no differences to controls by the third passage (P3 d11) (Figure 4D-E). In contrast, the 3x hypoxic group (P3 d11), showed a progressive increase in kidney injury and inflammatory markers including *LCN2, SAA1* and *SAA2*, markers associated with maladaptive repair (*S100A8* and *S100A9)* (52–54) and apoptosis (*CASP3*, *CASP9*) (Figure 4F-G). The compounding effect of repeated hypoxic injury was further reflected by the increase in proinflammatory and profibrotic mediators between P1 and P3 d11 injury groups including *IL1B* (fold change (FC,) = 2, adj. *p*-value = 0.01), *IL18* (FC = 1.6, adj. *p*-value = 0.01), *CXCL6* (FC = 3.6, adj. *p*-value < 0.001) and *CXCL5* (FC = 2.7, adj. *p*-value < 0.001) (Supplemental File 3). The transcriptional profile of normoxic organoids remained relatively consistent throughout the experiment though some epithelial markers like *CLU* and *UPK1B* were decreased in P3 (Supplemental Figure 4).

## Discussion

A variety of human organoid models are now available to study kidney development and disease (55). However, we lack data to evaluate the relative strengths and limitations of each model for applications in kidney development, physiology and disease. Here, we generated and compared tubuloids from adult nephrectomies, iPSC-derived kidney organoids and for the first time-human fetal kidneys. Comparison between the tubuloid models and benchmarking to whole iPSC-derived kidney organoids and adult kidneys revealed differences in cell type, regeneration, ageing, and repair signatures.

Reports of adult-derived tubuloids expressing nephron and collecting duct markers raised hope that they may represent more mature cell states than those in iPSC-derived kidney organoid (21, 22, 56). While select markers of the distal nephron and Loop of Henle were robustly expressed in tubuloids, the same was not observed for proximal tubule. iPSC-derived kidney organoids expressed higher levels of proximal tubule markers than any tubuloid culture, which challenges the proposition that adult tubuloids model the mature nephron epithelium. Why proximal tubule signatures are stronger in iPSC-derived kidney organoids than tubuloids remains unclear. Contributing factors may include suboptimal media conditions, dependence on cell communication with surrounding interstitial cells absent from tubuloid culture, or suppression of proximal identity by a de-differentiation and repair program activated with each passage. Despite limited marker expression, all tubuloid models retained cells with proximal identity and likely provide valuable opportunities to interrogate proximal tubule function and disorders. Optimizing media conditions to enhance the differentiation of specific nephron types is likely to improve epithelial cell identities in tubuloid cultures (57).

Collecting duct markers were strongly expressed in both adult and fetal tubuloids, with the latter showing a developing ureteric epithelium signature. These results suggest that the starting material influences the cell types and characteristics of the tubuloid culture. As such, tubuloids generated from different kidney regions likely vary in composition, and sampling the medulla may yield distinct collecting duct and nephron signatures.

Interexperimental variability in iPSC-derived organoids is well recognized (18). Differences in gene expression among biological replicates in all three tubuloid types shows that tubuloids are no exception. Nonetheless, conserved expression of multiple nephron segment markers in fetal, adult and iTubuloid cultures demonstrates their reproducible ability to model kidney epithelial cell types. Indeed, markers associated with nephron regeneration, cellular stress, inflammation and senescence were significantly increased in adult tubuloids, offering potential insights into nephron repair mechanisms and the effects of aging on kidney disease.

Long-term tubuloid cultures offer great potential for studying stressors that drive CKD progression. In this study, we showed that iTubuloids can mount a hypoxic response and effectively recover from a single injury, but multiple injuries compounded to induce signs of maladaptive repair. As such, tubuloids may provide a human platform to investigate the mechanisms of cumulative injury and screen for the mechanisms of maladaptive repair and potential therapies to prevent progression to CKD. Finally, our transcriptional profiling data provides a valuable resource to evaluate the suitability of adult tubuloids, fetal tubuloids and iTubuloid cultures for a range of exciting new opportunities in experimental and clinical nephrology.

## Acknowledgements

The authors acknowledge use of the MHTP Medical Genomics Facility, Monash Bioinformatics Platform and Monash Micro Imaging. This study used infrastructure enabled by Bioplatforms Australia and the National Collaborative Research Infrastructure Strategy at the Monash Proteomics and Metabolomics Platform. The authors thank Sara Howden and Melissa Little for support and providing the 522 and 808 iPSC lines, and Andrew Laslett for the D4C4 iPSC line

## Author Contributions

A.N.C., D.J.N.P. and A.B.N.N conceived and designed the study; A.B.N.N generated tubuloids, analysed data and generated figures; L.K and E.E provided nephrectomy samples; A.N.C., L.S., P.M. and E.D. collected or provided human fetal kidney tissue.

The original draft was written by A.B.N.N. and revised with A.N.C.; A.N.C. and D.J.N.P. supervised the project; All authors subsequently reviewed and approved the manuscript.

## Data Sharing Statement

Bulk RNAseq data has been deposited in the National Center for Biotechnology Information’s GEO database (accession No. GSE252055).

## References

1. Liu B-C, Tang T-T, Lv L-L. How Tubular Epithelial Cell Injury Contributes to Renal Fibrosis. In: Liu B-C, Lan H-Y, Lv L-L, editors. Renal Fibrosis: Mechanisms and Therapies. Singapore: Springer Singapore; 2019. p. 233–52.

2. Faria J, Ahmed S, Gerritsen KGF, Mihaila SM, Masereeuw R. Kidney-based in vitro models for drug-induced toxicity testing. Arch Toxicol. 2019;93(12):3397–418.

3. Digby JLM, Vanichapol T, Przepiorski A, Davidson AJ, Sander V. Evaluation of cisplatin-induced injury in human kidney organoids. Am J Physiol Renal Physiol. 2020;318(4):F971–f8.

4. Gupta N, Matsumoto T, Hiratsuka K, Garcia Saiz E, Galichon P, Miyoshi T, et al. Modeling injury and repair in kidney organoids reveals that homologous recombination governs tubular intrinsic repair. Sci Transl Med. 2022;14(634):eabj4772.

5. Schutgens F, Rookmaaker MB, Margaritis T, Rios A, Ammerlaan C, Jansen J, et al. Tubuloids derived from human adult kidney and urine for personalized disease modeling. Nat Biotechnol. 2019;37(3):303–13.

6. Przepiorski A, Vanichapol T, Espiritu EB, Crunk AE, Parasky E, McDaniels MD, et al. Modeling oxidative injury response in human kidney organoids. Stem Cell Res Ther. 2022;13(1):76.

7. Ohmori T, De S, Tanigawa S, Miike K, Islam M, Soga M, et al. Impaired NEPHRIN localization in kidney organoids derived from nephrotic patient iPS cells. Scientific Reports. 2021;11(1):3982.

8. Freedman BS, Brooks CR, Lam AQ, Fu H, Morizane R, Agrawal V, et al. Modelling kidney disease with CRISPR-mutant kidney organoids derived from human pluripotent epiblast spheroids. Nature Communications. 2015;6(1):8715.

9. Takasato M, Er PX, Chiu HS, Maier B, Baillie GJ, Ferguson C, et al. Kidney organoids from human iPS cells contain multiple lineages and model human nephrogenesis. Nature. 2015;526(7574):564–8.

10. Taguchi A, Kaku Y, Ohmori T, Sharmin S, Ogawa M, Sasaki H, et al. Redefining the in vivo origin of metanephric nephron progenitors enables generation of complex kidney structures from pluripotent stem cells. Cell Stem Cell. 2014;14(1):53–67.

11. Freedman BS, Brooks CR, Lam AQ, Fu H, Morizane R, Agrawal V, et al. Modelling kidney disease with CRISPR-mutant kidney organoids derived from human pluripotent epiblast spheroids. Nat Commun. 2015;6:8715.

12. Morizane R, Lam AQ, Freedman BS, Kishi S, Valerius MT, Bonventre JV. Nephron organoids derived from human pluripotent stem cells model kidney development and injury. Nat Biotechnol. 2015;33(11):1193–200.

13. Howden SE, Wilson SB, Groenewegen E, Starks L, Forbes TA, Tan KS, et al. Plasticity of distal nephron epithelia from human kidney organoids enables the induction of ureteric tip and stalk. Cell Stem Cell. 2021;28(4):671–84.e6.

14. Tanigawa S, Tanaka E, Miike K, Ohmori T, Inoue D, Cai C-L, et al. Generation of the organotypic kidney structure by integrating pluripotent stem cell-derived renal stroma. Nature Communications. 2022;13(1):611.

15. Combes AN, Zappia L, Er PX, Oshlack A, Little MH. Single-cell analysis reveals congruence between kidney organoids and human fetal kidney. Genome Medicine. 2019;11(1):3.

16. Little MH, Combes AN. Kidney organoids: accurate models or fortunate accidents. Genes Dev. 2019;33(19-20):1319–45.

17. Nunez-Nescolarde AB, Nikolic-Paterson DJ, Combes AN. Human Kidney Organoids and Tubuloids as Models of Complex Kidney Disease. The American Journal of Pathology. 2022.

18. Phipson B, Er PX, Combes AN, Forbes TA, Howden SE, Zappia L, et al. Evaluation of variability in human kidney organoids. Nat Methods. 2019;16(1):79–87.

19. Morizane R, Bonventre JV. Generation of nephron progenitor cells and kidney organoids from human pluripotent stem cells. Nature Protocols. 2017;12(1):195–207.

20. Subramanian A, Sidhom E-H, Emani M, Vernon K, Sahakian N, Zhou Y, et al. Single cell census of human kidney organoids shows reproducibility and diminished off-target cells after transplantation. Nature Communications. 2019;10(1):5462.

21. Schutgens F, Rookmaaker MB, Margaritis T, Rios A, Ammerlaan C, Jansen J, et al. Tubuloids derived from human adult kidney and urine for personalized disease modeling. Nature Biotechnology. 2019;37(3):303–13.

22. Gijzen L, Yousef Yengej FA, Schutgens F, Vormann MK, Ammerlaan CME, Nicolas A, et al. Culture and analysis of kidney tubuloids and perfused tubuloid cells-on-a-chip. Nat Protoc. 2021;16(4):2023–50.

23. Xu Y, Kuppe C, Perales-Patón J, Hayat S, Kranz J, Abdallah AT, et al. Adult human kidney organoids originate from CD24(+) cells and represent an advanced model for adult polycystic kidney disease. Nat Genet. 2022;54(11):1690–701.

24. Mashouf P, Tabibzadeh N, Kuraoka S, Oishi H, Morizane R. Cryopreservation of human kidney organoids. Cellular and Molecular Life Sciences. 2024;81(1):306.

25. Wiersma LE, Avramut MC, Lievers E, Rabelink TJ, van den Berg CW. Large-scale engineering of hiPSC-derived nephron sheets and cryopreservation of their progenitors. Stem Cell Research & Therapy. 2022;13(1):208.

26. Yousef Yengej FA, Jansen J, Ammerlaan CME, Dilmen E, Pou Casellas C, Masereeuw R, et al. Tubuloid culture enables long-term expansion of functional human kidney tubule epithelium from iPSC-derived organoids. Proc Natl Acad Sci U S A. 2023;120(6):e2216836120.

27. Nunez-Nescolarde AB, Piran M, Perlaza-Jimenez L, Barlow CK, Steele JR, Deveson D, et al. Hypoxic injury triggers maladaptive repair in human kidney organoids. bioRxiv. 2023:2023.10.04.558359.

28. Grubman A, Choo XY, Chew G, Ouyang JF, Sun G, Croft NP, et al. Transcriptional signature in microglia associated with Aβ plaque phagocytosis. Nature Communications. 2021;12(1):3015.

29. Perry A, Powell D. Laxy Genomics Pipelines. Zenodo. 2020.

30. Patel H, Ewels P, Peltzer A, Hammarén R, Botvinnik O, Sturm G, et al. nf-core/rnaseq: nf-core/rnaseq v3.2 - Copper Flamingo (3.2). Zenodo. 2021.

31. Patro R, Duggal G, Love MI, Irizarry RA, Kingsford C. Salmon provides fast and bias-aware quantification of transcript expression. Nat Methods. 2017;14(4):417–9.

32. Ewels P, Magnusson M, Lundin S, Käller M. MultiQC: summarize analysis results for multiple tools and samples in a single report. Bioinformatics. 2016;32(19):3047–8.

33. Powell D. Degust: interactive RNA-seq analysis. Drpowell/Degust. 2015;4(1):4.1.

34. Robinson MD, Oshlack A. A scaling normalization method for differential expression analysis of RNA-seq data. Genome Biology. 2010;11(3):R25.

35. Law CW, Chen Y, Shi W, Smyth GK. voom: Precision weights unlock linear model analysis tools for RNA-seq read counts. Genome biology. 2014;15(2):1–17.

36. Harrison PF. Varistran: Anscombe’s variance stabilizing transformation for RNA-seq gene expression data. Journal of Open Source Software. 2017;2(257).

37. Sherman BT, Hao M, Qiu J, Jiao X, Baseler MW, Lane HC, et al. DAVID: a web server for functional enrichment analysis and functional annotation of gene lists (2021 update). Nucleic acids research. 2022;50(W1):W216–W21.

38. Combes AN, Zappia L, Er PX, Oshlack A, Little MH. Single-cell analysis reveals congruence between kidney organoids and human fetal kidney. Genome Med. 2019;11(1):3.

39. Ransick A, Lindström NO, Liu J, Zhu Q, Guo JJ, Alvarado GF, et al. Single-Cell Profiling Reveals Sex, Lineage, and Regional Diversity in the Mouse Kidney. Dev Cell. 2019;51(3):399–413.e7.

40. Lindström NO, De Sena Brandine G, Tran T, Ransick A, Suh G, Guo J, et al. Progressive Recruitment of Mesenchymal Progenitors Reveals a Time-Dependent Process of Cell Fate Acquisition in Mouse and Human Nephrogenesis. Dev Cell. 2018;45(5):651–60.e4.

41. Wu H, Malone AF, Donnelly EL, Kirita Y, Uchimura K, Ramakrishnan SM, et al. Single-Cell Transcriptomics of a Human Kidney Allograft Biopsy Specimen Defines a Diverse Inflammatory Response. J Am Soc Nephrol. 2018;29(8):2069–80.

42. Wu H, Uchimura K, Donnelly EL, Kirita Y, Morris SA, Humphreys BD. Comparative Analysis and Refinement of Human PSC-Derived Kidney Organoid Differentiation with Single-Cell Transcriptomics. Cell Stem Cell. 2018;23(6):869–81.e8.

43. Gerli MFM, Calà G, Beesley MA, Sina B, Tullie L, Sun KY, et al. Single-cell guided prenatal derivation of primary fetal epithelial organoids from human amniotic and tracheal fluids. Nature Medicine. 2024;30(3):875–87.

44. de Magalhães JP, Curado J, Church GM. Meta-analysis of age-related gene expression profiles identifies common signatures of aging. Bioinformatics. 2009;25(7):875–81.

45. Takemon Y, Chick JM, Gerdes Gyuricza I, Skelly DA, Devuyst O, Gygi SP, et al. Proteomic and transcriptomic profiling reveal different aspects of aging in the kidney. eLife. 2021;10:e62585.

46. Rodwell GE, Sonu R, Zahn JM, Lund J, Wilhelmy J, Wang L, et al. A transcriptional profile of aging in the human kidney. PLoS Biol. 2004;2(12):e427.

47. Combes AN, Phipson B, Lawlor KT, Dorison A, Patrick R, Zappia L, et al. Single cell analysis of the developing mouse kidney provides deeper insight into marker gene expression and ligand-receptor crosstalk. Development. 2019;146(12).

48. Kolvenbach CM, Shril S, Hildebrandt F. The genetics and pathogenesis of CAKUT. Nature Reviews Nephrology. 2023;19(11):709–20.

49. Siew ED, Parr SK, Abdel-Kader K, Eden SK, Peterson JF, Bansal N, et al. Predictors of Recurrent AKI. Journal of the American Society of Nephrology. 2016;27(4).

50. Chen P-S, Chiu W-T, Hsu P-L, Lin S-C, Peng IC, Wang C-Y, et al. Pathophysiological implications of hypoxia in human diseases. Journal of Biomedical Science. 2020;27(1):63.

51. Xu L. The Role of Myeloid Cells in Acute Kidney Injury and Kidney Repair. Kidney360. 2021;2(11):1852–64.

52. Balzer MS, Doke T, Yang YW, Aldridge DL, Hu H, Mai H, et al. Single-cell analysis highlights differences in druggable pathways underlying adaptive or fibrotic kidney regeneration. Nat Commun. 2022;13(1):4018.

53. Gerhardt LMS, Liu J, Koppitch K, Cippà PE, McMahon AP. Single-nuclear transcriptomics reveals diversity of proximal tubule cell states in a dynamic response to acute kidney injury. Proceedings of the National Academy of Sciences. 2021;118(27):e2026684118.

54. Zhang F, Zhou X, Zou H, Liu L, Li X, Ruan Y, et al. SAA1 is transcriptionally activated by STAT3 and accelerates renal interstitial fibrosis by inducing endoplasmic reticulum stress. Exp Cell Res. 2021;408(1):112856.

55. Nunez-Nescolarde AB, Nikolic-Paterson DJ, Combes AN. Human Kidney Organoids and Tubuloids as Models of Complex Kidney Disease. The American Journal of Pathology. 2022;192(5):738–49.

56. Yousef Yengej FA, Jansen J, Rookmaaker MB, Verhaar MC, Clevers H. Kidney Organoids and Tubuloids. Cells. 2020;9(6):1326.

57. Schnell J, Achieng M, Lindstrom NO. Principles of human and mouse nephron development. Nat Rev Nephrol. 2022;18(10):628–42.

