## Supplemental for "Comparative analysis of iPSC-derived human kidney organoid and tubuloid models"

Nunez-Nescolarde A.B., et al

The PDF file includes:
Materials and Methods
Supplemental Figures 1 to 4
Supplemental Tables 1 and 2

Other Supplemental Material for this manuscript includes the following:

Supplemental Files S1-3 and Video S1 and S2

### Materials and Methods

#### iPSC lines and human material

The three **iPSC lines** in this study were used under material transfer agreements with the Commonwealth Scientific and Industrial Research Organization (CSIRO, 808 line) and Murdoch Children's Research Institute (MCRI, 522 and D4C4 lines). Sex chromosomes were defined as follows: 808 XY, 522 XX, D4C4 XX. **Nephrectomy samples** were obtained with informed patient consent and approval from the Austin Hospital Human Research Ethics Committee (HUREC) (#22635/LNR/13/Austin/163). Radical nephrectomies were performed for renal cell carcinoma. Adjacent healthy tissue was identified and collected from the renal cortex by an expert pathologist. Nephrectomy samples and related data were provided by the Victorian Cancer Biobank, supported by the Victorian Government, Australia (donor 1 – XX, donor 2 and 3 – XY). **Human fetal tissues** were obtained with informed patient consent from the Royal Women's Hospital (RWH) and approval from HURECs at RWH (#EMR RMH 95016) and Monash University (#37642). Fetal sample IDs, age in weeks of gestation (w) and sex: 369 14w XX, 389 12w XX, 399 13w XX.

#### iPSC Culture

Cells were maintained in Matrigel-coated (Corning, cat no. FAL35427) 6-well plates in essential 8 (E8) medium (Thermo Fisher Scientific) supplemented with 1% Penicillin/Streptomycin (Pen/Strep) (Thermo Fisher Scientific). Cells were passaged every 3-4 days once wells reached 70-80% confluency and split 1:3 using 0.5 mM EDTA (Thermo Fisher Scientific).

#### Repetitive hypoxic Injury

After confirming robustness of the iTubuloid model using three independent iPSC lines, we used iTubuloids derived from the 522.3 iPSC line for the repetitive injury experiment. Passage 7 iTubuloids were divided in three groups: control, 1 time hypoxic injury (1x hypoxia), and 3 times hypoxic injury (3x hypoxia). Control groups were maintained in normoxic (19% $O_2$) conditions during the experiment. The hypoxic exposure was performed 4 days after each passage for a total of 48h (incubators set at 1% $O_2$ levels). Wells with iTubuloids were harvested for analysis at day 6 or at day 11, after a 5-day recovery period in normoxic conditions. Three wells (n=3) were harvested for each group. For each passage, a total of 4 wells were pooled during each passage to generate 10 new wells. For bulk RNAseq wells were snap-frozen.

**Immunostaining**

Tubuloids were cultured in glass-bottom dishes (150680, Themor Fisher), with Matrigel droplets of 30 μl. Tubuloids were fixed in 4% formaldehyde for 16 hr at 4°C with constant rolling, followed by 20 min blocking solution (5 % Donkey serum, 0.1 % Triton X-100, made up in PBS) at room temperature. Tubuloids were then placed overnight with the primary antibody at 4 °C. After 30 min wash with PBS, tubuloids were incubated with secondary antibody for 2 hours at room temperature with constant rolling. After washing for 20 min with PBS, a final wash was performed with Milli-Q-purified water to prevent crystal formation. The list of antibodies used can be found in Table S2. Fluorescence images were collected using the 3i CSU-W1 T2 Yokogawa Spinning disk confocal. Representative images were generated in FIJI.

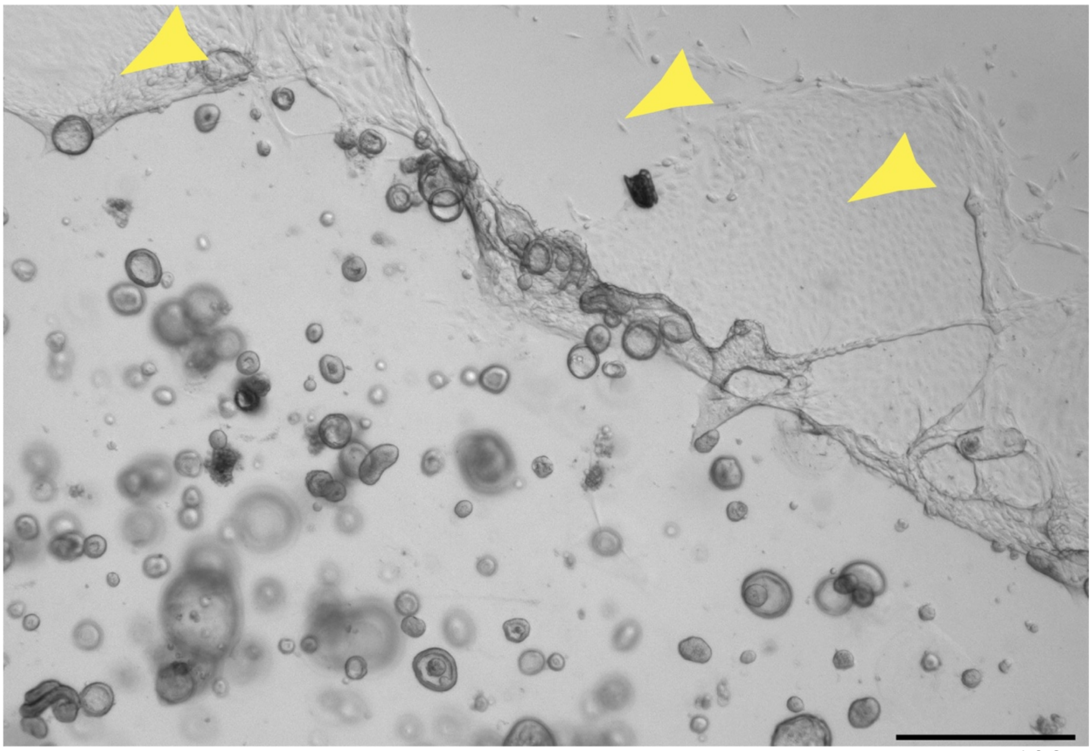

**Supplemental Figure 1. Bright field image of fibroblast/stroma contamination on tubuloid cultures.** Well of iTubuloid at day 3 after passage number 9. Yellow arrow showing cells with fibroblast/stroma morphology.

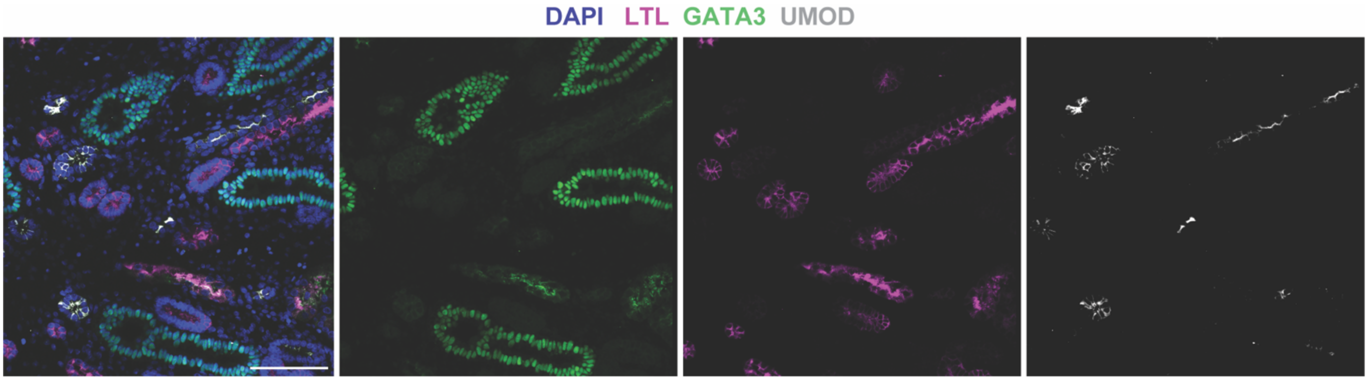


**Supplemental Figure 2. Human fetal kidney section showing complete separation of proximal and distal markers.** Representative immunofluorescence images showing LTL (proximal tubule marker), GATA3 (distal marker) and UMOD (loop of Henle). Scale bars = 100 µm.

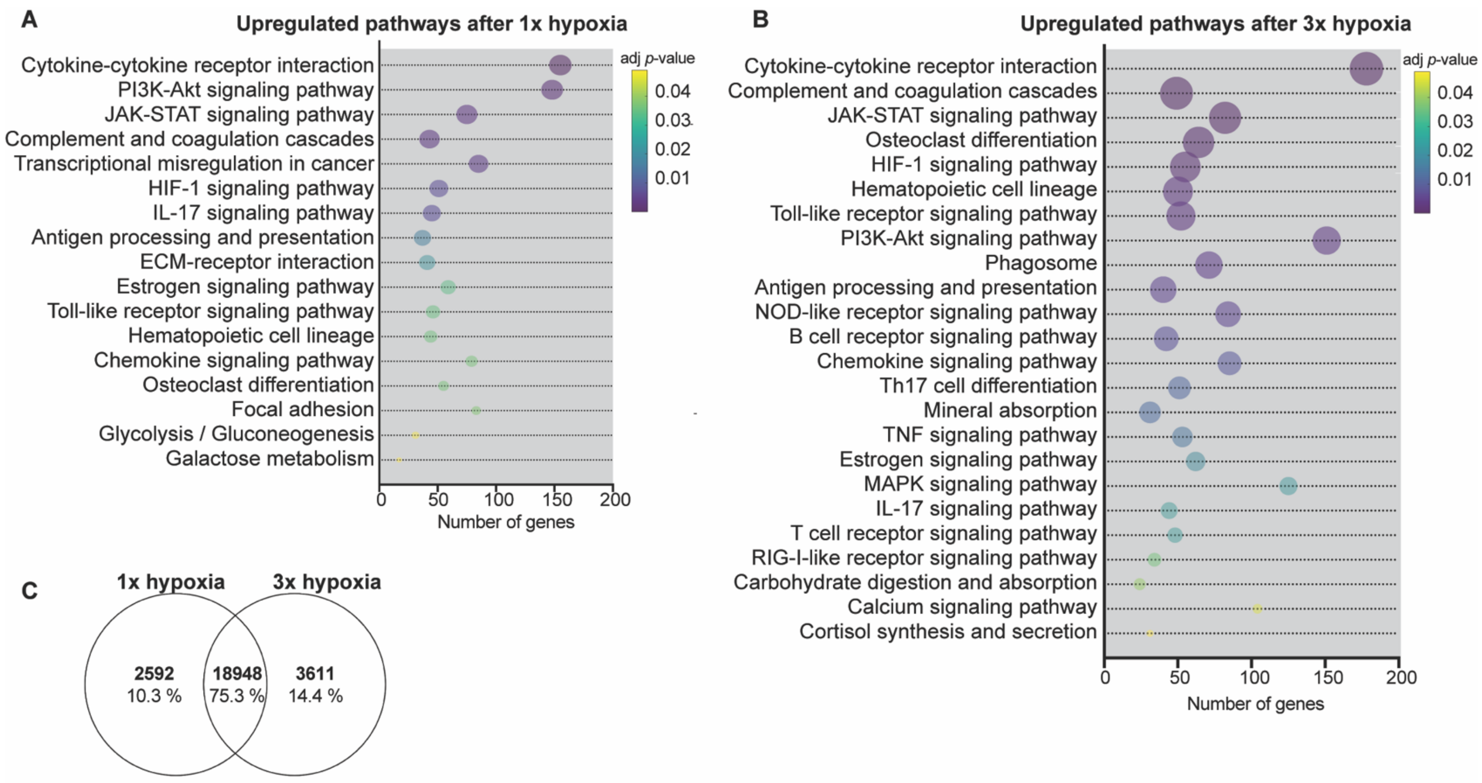


**Supplemental Figure 3. Successful recovery after 1x hypoxic exposure.** (A) KEGG pathways upregulated in 1x hypoxic exposed tubuloids compared to controls (P1 d6). (B) KEGG pathways upregulated in 3x hypoxic exposed tubuloids compared to stage-matched controls (P3 d6). Enrichment analysis for A and B performed with DAVID software, using genes with $\log_2$ fold change > 0.5 and *p*-value < 0.05. Ball size indicates the number of genes found in each pathway. (C) Venn diagram illustrating unique and common differentially expressed genes in 1x and 3x injured iTubuloids at day 6.

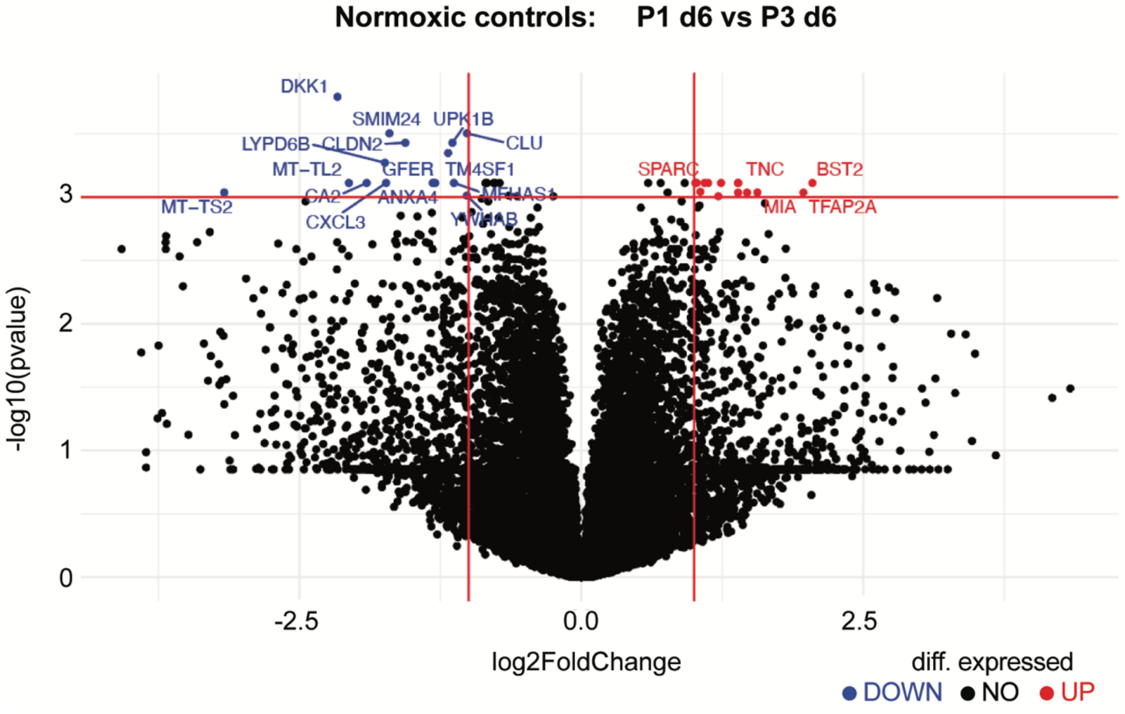


**Supplemental Figure 4. iTubuloid culture remain stable over time.** Volcano plot of differentially expressed genes between P1 and P3 normoxic controls, both at d6. Genes that are downregulated are depicted in blue, upregulated in red and no differentially expressed in black. Differentially expressed genes were established based on a log2 Fold Change > 1 and adjusted p-value < 0.001 (Benjamini-Hochberg method).

**Table S1.** Reagents

| Name | Supplier | Manufacturer | Catalogue number |
|---|---|---|---|
| Advanced DMEM/F12 | Thermo Fisher | Gibco | 12634010 |
| HEPES | Thermo Fisher | Gibco | 15630080 |
| GlutaMax | Thermo Fisher | Gibco | 35050061 |
| Penicillin-streptomycin | Thermo Fisher | Gibco | 15140122 |
| R-spondin-3-conditioned medium | Produced by the Monash BDI organoid Program | | |
| Recombinant human EGF | Lonza | Peprotech | AF-100-15 |
| N-Acetyl-L-cysteine (NAC) | Merck | Sigma-Aldrich | A9165 |
| A 83-01 | In vitro technologies | Tocris | Tocris A-83-01 (RDS293910) |
| Primocin | Integrated Sciences | InvivoGen | ant-pm-2 |
| Matrigel | In vitro technologies | Corning | 356231 |
| Collagenase A | MP Biomedicals | Sigma-Aldrich | 10103578001 |
| B27 supplement | Thermo Fisher | Gibco | 17504044 |
| Recombinant human FGF-10 | Lonza | Peprotech | 100-26-500 |
| Recombinant human FGF-2 | Lonza | Peprotech | 100-18B-500 |
| TrypLE Select Enzyme (1X) | Thermo Fisher | Gibco | 12563011 |
| Y-27632 (dihydrochloride) | In vitro technologies | Tocris | 1254 |

**Table S2.** Antibody list.

| Primary | | | | |
|---|---|---|---|---|
| **Target** | **Species** | **Concentration** | **Catalogue number** | **Company** |
| GATA3 | Mouse | 1:300 | MA1-028 | Life Technologies |
| LTL | Biotin | 1:400 | B-1325 | Vector Technologies |
| CK | Mouse | 1:200 | ab115959 | Abcam |
| MKI67 | Biotin | 1:300 | 13-5698-82 | Invitrogen |

| | | | | |
|---|---|---|---|---|
| SOX9 | Rabbit | 1:300 | Given by Dagmar Wilhelm | |
| **Secondary** | | | | |
| **Name** | | **Concentration** | **Catalogue number** | **Company** |
| Donkey anti-Mouse IgG | | 1:1000 | A32787 | Thermo Fisher Scientific |
| Alexa Fluor 568 streptavidin | | 1:1000 | S11226 | Thermo Fisher Scientific |
| Donkey anti-Rabbit IgG | | 1:1000 | A1004200 | Thermo Fisher Scientific |
